# Characterization and Phylogenetic Insights into the First Viral Major Vault Proteins (MVPs), Identified in Bacteriophages from the Human Gut

**DOI:** 10.1101/2023.11.10.566534

**Authors:** Thiago M. Santos, Emerson Galvão, Helena Gomes, Luiz-Eduardo Del-Bem

## Abstract

The Major Vault Protein (MVP) is the principal component of the enigmatic vault complex, dome-shaped cytoplasmic ribonucleoproteins found in eukaryotes. Although vaults have been associated with several minor functions, including amino acid storage and cargo transport, their overall function remains to be fully understood. While MVPs are widely distributed among most higher eukaryotes and have homologs in prokaryotes, no similar protein have been reported in viral species until now. In this study, we explored the NCBI viral protein database for MVP homologs and identified two putative MVPs from distinct tailed bacteriophage isolates from the Class *Caudoviricetes*, ctOa11 and ctRi15, of human gut metagenomic samples. Further *in silico* analysis were conducted to describe and characterize these unprecedent phage proteins. Phylogenetic topologies indicate that viral MVPs are closely related to bacterial MVP homologs. Tridimensional protein models of viral MVPs were generated to perform protein structure comparisons with eukaryotic and prokaryotic MVPs, indicating a significant similarity to bacterial proteins. For the functional significance of MVPs in phage infectivity, we suggest that they may have an unknown metabolic role enhancing phage fitness. While this work provides new insights into the genetic phage diversity and phylogenetic distribution of MVPs, complementary functional studies are necessary to fully elucidate their function in viral and bacterial species.

## 1. INTRODUCTION

Vaults are the largest known eukaryotic ribonucleoprotein complexes described to date, yet studies involving this structure are scarce. Mammalian vaults are 13 MDa dome-shaped complexes, mostly composed by multiple copies of the Major Vault Protein (MVP) and by a minor proportion of vault Poly-ADP Ribose Polymerase (vPARP), Telomerase-associated Protein 1 (TEP1) and vault RNA (vRNA) (Kedersha et al., 1990; Kedersha & Rome, 1986; Kickhoefer, Siva, et al., 1999, p. 193; Kickhoefer, Stephen, et al., 1999). vPARP belongs to a family of enzymes that catalyze the attachment of poly-ADP-ribose to other proteins, including itself, and is mainly involved in processes related to genomic stability (Kickhoefer, Siva, et al., 1999). This protein was also shown to ribosylate MVPs (Kickhoefer, Siva, et al., 1999) and is required for Vault-mediated drug resistance in ovarian cancer cells lines (Wojtowicz, Karolina et al., 2017), however, its ultimate function in vaults remains elusive. TEP1, an RNA binding protein, is part of the telomerase complex, responsible for telomere maintenance (Harrington et al., 1997) and, in vaults, it is believed to stabilize the integration of vRNAs into its structure (Kickhoefer et al., 2001; Poderycki et al., 2005). vRNAs are highly conserved non-coding RNAs of 88 to 140 nucleotides that can be associated with the vault structure (Kickhoefer et al., 1993). The role of vRNAs is still under investigation, but suggested functions involve drug resistance, cell proliferation, apoptosis and nutrient sequestering (Hahne et al., 2021; SHAIK, 2013).

Undoubtedly, however, is the essentiality of the MVP for the formation of the vault’s structure. Composing the shell of the ribonucleoprotein complex, this protein constitutes over 70% of the vault’s mass (Rome et al., 1991). MVPs spontaneously oligomerize to form the vault structure (**Fig. 1**) (Stephen et al., 2001). About 78 MVPs are necessary to assemble a complete vault, which consists of two halves of 39 MVPs each interacting with each other through non-covalent interactions at the vault waist region (Kato et al., 2008). The MVP typically consists of a multiple N-terminal repeat domain, a shoulder domain, a cap-helix domain, and a highly disordered cap-ring region at the C-terminal end. The shoulder domain connects the repeat domain to the cap-helix, a 42-turn a-helical domain. The cap-helix domain is responsible for self-assembly through lateral oligomerization of MVP proteins (Tanaka et al., 2009; van Zon et al., 2002).

**Fig. 1.**
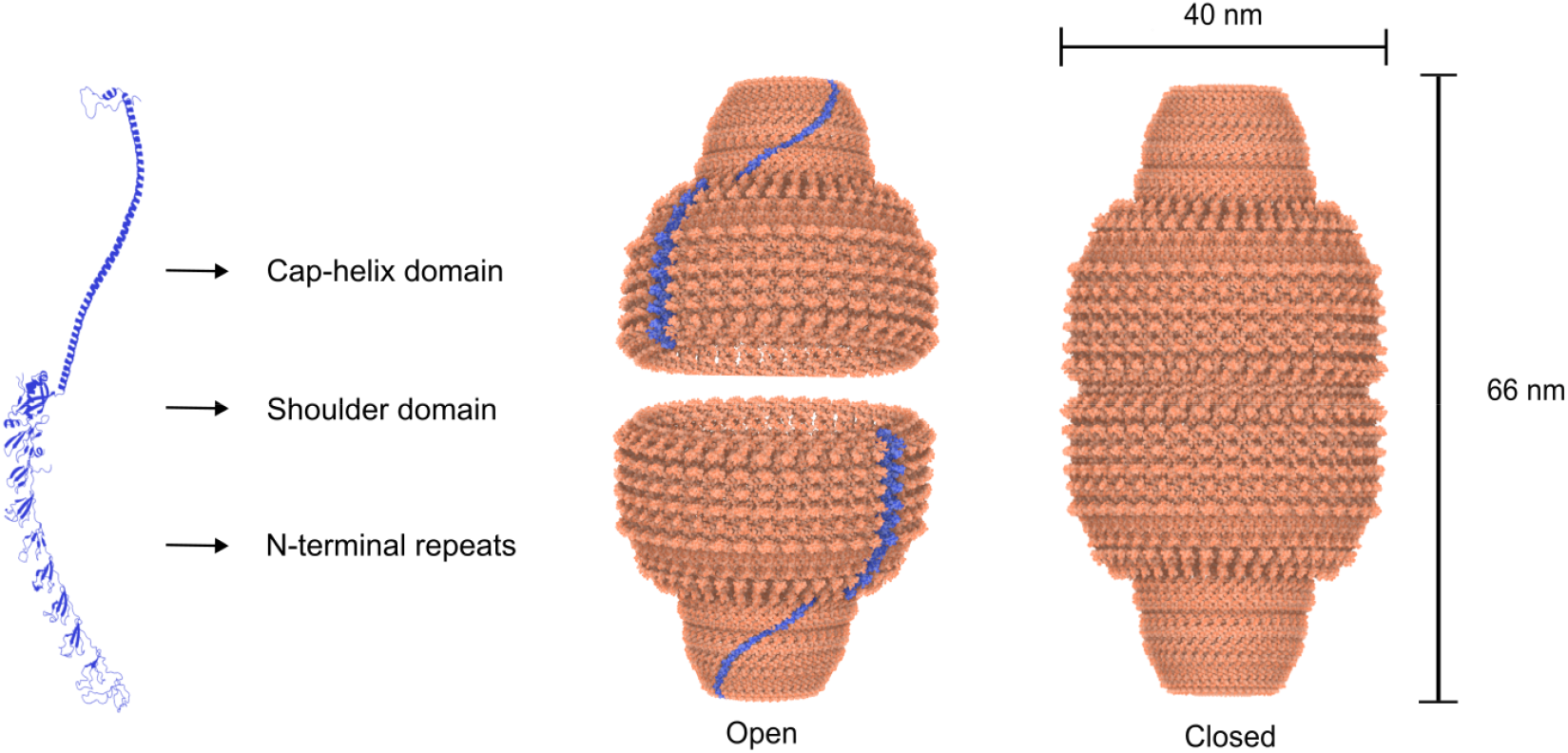
Major Vault Protein (MVP) and Vault structure. MVP monomer is composed by N-terminal repeats, consisting of 9 MVP domains; the shoulder domain, which connects the Cap-helix domain to the N-terminal repeats; and the Cap-helix domain, a long alpha-helix segment. MVPs oligomerize to form a half vault. Two halves are linked by the waist region to form a complete vault. Shown here are MVP and Vault structures from rat liver at 3.5 angstrom resolution (PDB-4V60).

Although the precise role of vaults is not yet fully understood, extensive efforts have been made to investigate their functions. Experimental studies in mammalian and yeast cells suggest that the functions of vaults can be broadly categorized into a few areas. First, vaults may participate in several signaling pathways, likely through the nucleocytoplasmic shuttle of molecules, such as estrogen receptors and Phosphatase and TENsin homolog (PTEN), a tumor suppressor protein (Abbondanza et al., 1998; Yu et al., 2002). Second, there is considerable focus on the multidrug resistance effect of vaults. In fact, MVP was initially labeled as Lung Resistance-related Protein (LRP) due to its first documented phenotypic association with lung cancer (Scheffer et al., 1995). Overexpression of MVPs has been demonstrated in numerous cancer studies, indicating an important role in chemotherapy resistance during cancer treatment (List et al., 1996; Mossink et al., 2003; Ramberg et al., 2021; Scheffer et al., 1995, 2000). Additionally, vaults seem to participate in immune responses to viral infection. Studies have shown that *MVP* is an interferon-induced gene and is overexpressed during many viral infections (Wang et al., 2020). Finally, an interesting hypothesis proposes that vaults may be involved in nutrition stress management, functioning as storage devices of amino acids and nucleotides for later use during adverse conditions. Reduced survival rates of *Dictyostelium MVP*-/- under stressful nutrient conditions provide evidence for this hypothesis (SHAIK, 2013; Vasu & Rome, 1995).

In terms of phylogenetic distribution, MVP can be found in a wide range of eukaryotes forming highly conserved protein structures. Vaults of similar size and structure have been isolated from invertebrates such as sea urchin and the amoebozoan slime mold *Dictyostelium* (Hamill & Suprenant, 1997; Kedersha et al., 1990), suggesting important functions across eukaryotes. However, they are absent from plants and model invertebrates such as *Drosophila melanogaster* and *Caenorhabditis elegans* (SHAIK, 2013). Computational and experimental investigations have previously established the presence of *MVP* homologs in Cyanobacteria and bacteria from the Orders Myxococcales, Cytophagales, and Oscillatoriales (SHAIK, 2013; Sokolskyi, 2019). Notably, however, no *MVP* gene has ever been detected in viral genomes. Herein, we report and explore for the first time the presence of *MVP* genes in bacteriophages, from a poorly characterized lineage, sourced from the human gut microbiome.

## 2. METHODS

### 2.1 Sequence similarity search

To identify potential MVP homologs in viral protein databases, we utilized the amino acid sequence of the human MVP, a *bona-fide* model protein (UniProt accession number: Q14764), as a query in a PSI-BLAST (Altschul et al., 1997) search of the the Non-Redundant (NR) database at NCBI, specifically focusing on viral proteins, employing the BLOSUM62 matrix with an e-value threshold of 1.10^-05^. Two viral hits were obtained from different bacteriophages, which were then used as queries for a second round of similarity search against the NR database under the same parameters. Top hits were kept for further analysis. Another BLASTp search was conducted against the complete proteome of the two MVP-like bearing phages to determine the co-presence of TEP1 (UniProt accession number: Q99973) and vPARP (UniProt accession number: Q9UKK3) homologs using the human proteins as queries.

### 2.2 Phage Genomic Characterization and Taxonomy

To assess genome assembly quality and read coverage, sequence reads from which the two phages were derived (SRR1804492, SRR5156069, SRR6028456) were downloaded from the Sequence Read Archive (SRA) and mapped onto their deposited genomes using Bowtie2 v2.4.5 (Langmead & Salzberg, 2012). To express viral abundance, Reads Per Kilobase Million (RPKM) values were calculated for the entire genomes. Read alignment to the reference genomes was visualized using the IGV tool (Robinson et al., 2011). Circular genome maps of each viral species were generated using Proksee (Stothard et al., 2019).

To evaluate the conservation of gene order in these two phage genomes, a BLASTn search was performed utilizing less stringent parameters (word size 7 and with Filters and Masking deactivated). Also, to verify the gene order of the *MVP* flanking genes, a BLASTp search was conducted using 10 proteins upstream and 10 proteins downstream of the *MVP* gene from both phage genomes, with the same parameters previously applied.

The phage lifestyle prediction tools PhaTYP (Shang et al., 2023), PHACTS (McNair et al., 2012) and BACPHLIP (Hockenberry & Wilke, 2021) were applied to ctOa11 and ct2Ri15 to predict whether the viruses have a temperate lifestyle by integrating themselves into the bacterial genome, or a lytic lifestyle, involving the destruction of the bacterial cell. PhaTYP and BACPHLIP use genomic data, while PHACTS relies on the proteome.

To enhance viral taxonomy, the genomes of both phages were submitted to VipTree, a server designed for proteome-based phylogeny and the classification of viruses, also capable of predicting the most probable host candidates. This tool employs comprehensive genome-wide similarities estimated by tBLASTx, utilizing the Virus-Host DB database (RefSeq release 219). Additionally, Amino acid Average Identity (AAI) analysis was performed using the AAI calculator (Rodriguez-R & Konstantinidis, 2014). This method estimates the average amino acid identity based on the best hits and reciprocal best hits between two proteome files. AAI analysis provides a useful metric for comparing genome-wide protein content similarity between two or more genomes and can provide insights into evolutionary relationships among organisms.

### 2.3 Phylogenetic analysis of MVPs and MVP-like proteins

Amino acid sequences of the potential phage MVPs, their bacterial homologs and eukaryotic MVPs obtained from the UniProt database (98 sequences total) were used to reconstruct a phylogenetic tree to represent the evolutionary relationships among them. First, the sequences were aligned using MAFFT v7.453 (Katoh et al., 2002) applying the L-INS-I refinement method and BLOSUM62 matrix. The alignment was edited using the CIAlign software (Tumescheit et al., 2022) to remove gap-only columns, insertions and low-quality ends. The final alignment was next used as an input for the IQ-TREE 1.6.12 software (Nguyen et al., 2015) to perform the phylogenetic tree reconstruction. Model finder (Kalyaanamoorthy et al., 2017) estimated LG+I+G4 as the most suitable amino acid substitution model. The initial consensus tree was obtained by Maximum Likelihood and branch support was calculated by aLRT SH-like (Anisimova & Gascuel, 2006) in addition to Ultrafast Bootstrap method (Minh et al., 2013). Final trees were visualized and edited using FigTree v.1.4.4 (http://tree.bio.ed.ac.uk/software/figtree/).

### 2.4 Protein structural analyses

Based on amino acid sequences, tridimensional protein structures of putative phage and bacterial MVPs were modelled using ColabFold, an implementation of AlfaFold2 protein fold prediction system, in Google Colaboratory (Mirdita et al., 2022). Top ranked structures, based on local-distance difference test (pLDDT) and Predicted Aligned Error (PAE), were selected for each putative MVP. Human MVP was downloaded from the AlphaFold protein structure database (Varadi et al., 2022) and used as an eukaryotic reference model.

The predicted models were then submitted to PyMol v2.5.2 (The PyMOL Molecular Graphics System, Schrödinger, LLC) for protein structure comparison. Root Mean Square Deviation (RMSD) values were calculated for each protein model using an all-against-all alignment scheme. Protein images were created using the 3D Protein Imaging server (Tomasello et al., 2020).

## 3. RESULTS

### 3.1 MVP sequence similarity search and prophage insights

A BLASTp search was conducted in the NCBI virus protein database using the human MVP as a query, which unexpectedly revealed two positive hits (DAN98122.1, DAT09870.1) from the metagenome-assembled genomes (MAGs) of viral isolates ctOa11 (BK048656.1) and ct2Ri15 (BK042946.1), respectively (**Fig. 2A**). Both proteins were already automatically annotated by NCBI as putative MVPs, and the viruses were taxonomically assigned to the Class *Caudoviricetes*, a group of bacteriophages with head-tail morphology. These proteins were predicted by NCBI from metagenomic assemblies of human gut samples deposited in the SRA. The phage proteins showed an identity of about 22.8% when compared to the human MVP, with a shared identity of 49% among themselves. No positive hits were found for *TEP1* and *vPARP* in the phage genomes.

**Fig. 2.**
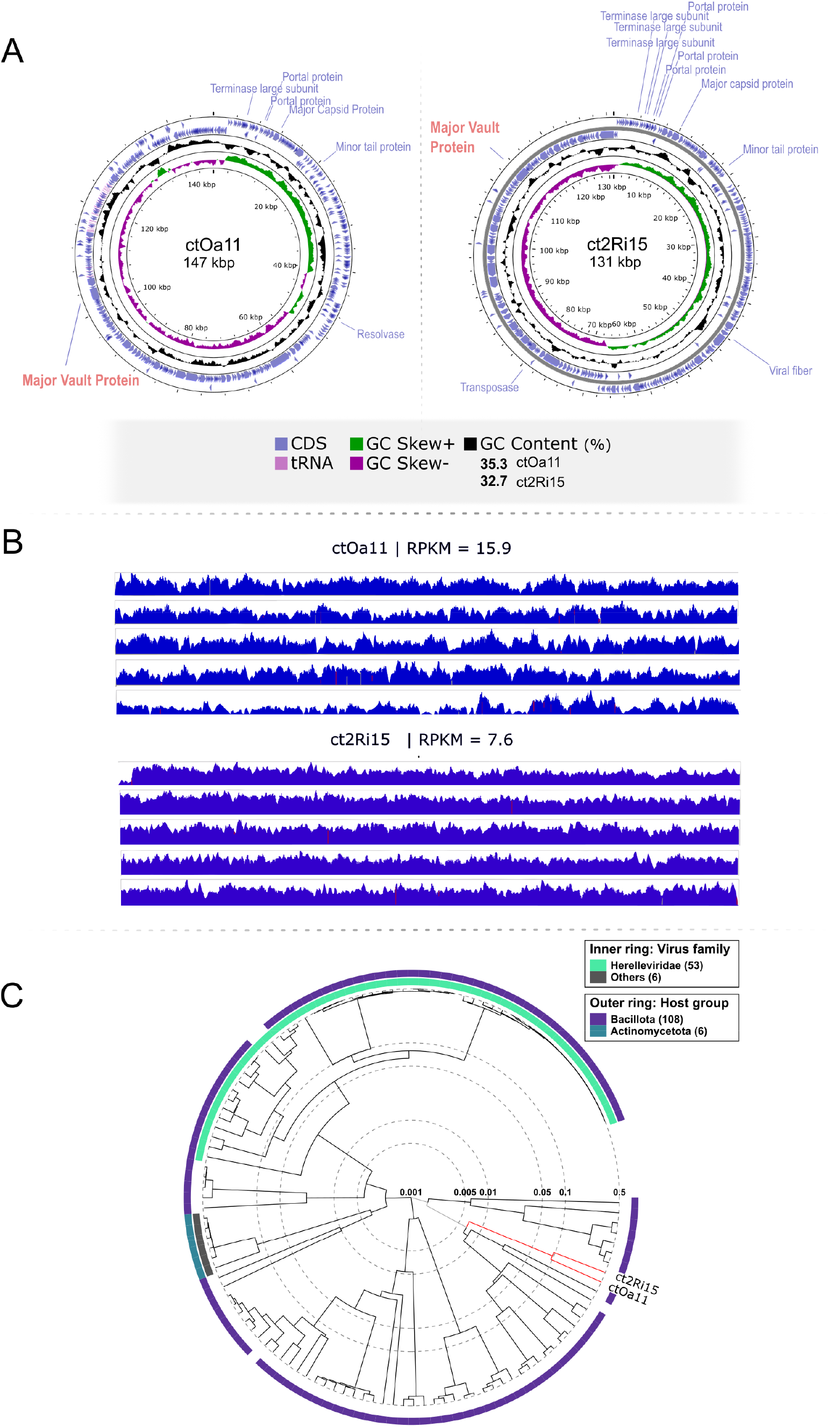
Viral genome analysis and taxonomy of *Caudoviricetes* sp. ct2Ri15 and ctOa11 **A)** Circular genome map of metagenomic-assembled bacteriophages (Most protein names are omitted for best visualization). Highlighted are the locations of the Major Vault Protein and the structural clusters **B)** Read coverage of assembled genomes of the phages and their respective RPKM values **C)** Proteomic tree generated by VipTree based on genome-wide similarities as determined by tBLASTx plotted on a log scale. Right and left lines are color-coded according to host group and virus family, respectively. For reference, ct2Ri15 and ctOa11 phages are highlighted in red. ct2Ri15 and ctOa11 could not be assigned to existing viral families and grouped together with other unassigned bacteriophages. A more detailed tree can be seen in **Supplemental Figure S2**. RPKM, Reads Per Kilobase Million.

The phage MVPs homologs were further used as query for a general search in the NR database, resulting in the discovery of significantly similar proteins. This step yielded prokaryotic hits only, with an average of 43% percentage identity, and stronger e-values. One interesting hit was MCI7444589.1, which is a hypothetical protein from the *Clostridium* sp. isolate SUG910 (GCA_022785265.1), derived from pig gut metagenome. This protein shares a 99.01% identity with a 100% protein coverage with the MVP from isolate ct2Ri15.

Due the high level of protein sequence similarity, we conducted a prophage search on the genomic data of *Clostridium* sp. isolate SUG910 using PHASTER, a server for the identification and annotation of prophage sequences within bacterial genomes (Arndt et al., 2016). No complete genome or assembly was available for this bacterial isolate; only contigs from the Whole Genome Shotgun (WGS) project where it was derived were at our disposal.

Three positive incomplete regions of prophages were predicted within the contig where the bacterial *MVP* gene was identified (**Supplemental Figure S1A**). One of these regions, region 2 (65859-76739bp), encompasses a 10.8kb segment populated with predicted 8 phage hit proteins. These proteins are associated with phages that infect species of *Bacillus*, *Streptomyces*, *Mycobacterium*, *Campylobacter,* and other bacterial species. The bacterial *MVP* gene is located immediately following this region (77006-79444bp).

To complement these findings, we investigated the genetic homology between ct2Ri15 and the *Clostridium* sp. contig using BLASTn and BLASTp (default parameters). The results revealed that 83% of the phage genome, with a 94.98% identity, is homologous to the *Clostridium* sp. contig (**Supplemental Figure S1B**). A BLASTp search, focusing on the region flanking the *MVP* genes, showed that one protein upstream and two proteins downstream of the bacterial *MVP* gene share over 95% similarity with their phage counterparts, while the subsequent proteins do not exhibit homology.

### 3.2 Phage Genomic Characterization and Taxonomy

DNA sequence reads obtained from the metagenomic samples where viral isolates were identified, were accurately mapped by Bowtie2 on the assembled genomes of ctOa11 and ct2Ri15 (**Fig. 2B**). No gaps were observed at the limits of the *MVP* gene, and sufficing read coverage was observed for the entire genome sequence of the phages. RPKMs of both whole genomes were computed to be 15.9 for ctOa11 and 7.6 for ct2Ri15.

PhaTYP (Shang et al., 2023), PHACTS (McNair et al., 2012), and BACPHLIP (Hockenberry & Wilke, 2021) predicted that phage ct2Ri15 has a temperate, lytic, and lytic lifestyle, respectively. Conversely, all three tools concur that phage ctOa11 exhibits a lytic lifestyle.

The phylogenetic tree built by VipTree was unable to assign ctOa11 and ct2Ri15 to existing viral families. Phages in the same clade lack family, genus, or species information. Concerning the host candidates, VipTree could not assign hosts to the phages either, however, other phages in the same clade infects bacteria from the phylum *Baciliota* (*Firmicutes*). Both phage genomes cluster together in this tree, branching out from the same node (**Fig. 2C, Supplemental Figure S2**) suggesting proximity. Since ct2Ri15 was assembled from metagenomic samples obtained from crAssphage amplicons (SRA Sample SRR1804492), there is a possibility that this phage could be a crAssphage-type phage. To investigate this, we conducted a whole-genome BLASTn search of both genomes as queries against the *Crassvirales* NCBI database and no significant hits were obtained.

Partial conservation of gene order becomes evident when comparing both whole phage genomes. Nucleotide similarity persists in approximately 44kbp regions of both genomes (**Supplemental Figure S3**). However, in terms of gene product order in the vicinity of the *MVP* gene in both phage genomes, distinct genes are observed in each region, indicating potential genomic rearrangements or divergent evolution. The AAI calculation estimates a 47.56% amino acid identity between the two phage proteomes.

### 3.3 Phylogenetic analysis of MVPs and MVP-like proteins

To visualize the evolutionary dynamics of the viral MVP homologs, along with the prokaryotic best-hits and eukaryotic MVPs, we built maximum likelihood phylogenetic trees. The reconstruction yielded a tree with the viral MVP homologs positioned within a bacterial subclade (**Fig. 3A, Supplemental Figure S4**), suggesting a degree of protein similarity between these two groups. The eukaryotic proteins clustered together after a long branch, implying a significant evolutionary distance between these clades. Within this cluster, cyanobacterial (*Oscillatoria acuminata*, *Oxynema aestuarii*, *Allocoleopsis franciscana*) MVPs are positioned at the base, while Myxococales (*Corallococcus coralloides* and *Melittangium boletus*) MVPs emerge as a sister group. Furthermore, looking at possible differences in protein length, eukaryotes present a higher average length of MVPs (858.78 AA) compared with the bacterial MVPs (822.43 AA) and viral MVPs (811.5 AA) (**Fig. 3B**).

**Fig. 3.**
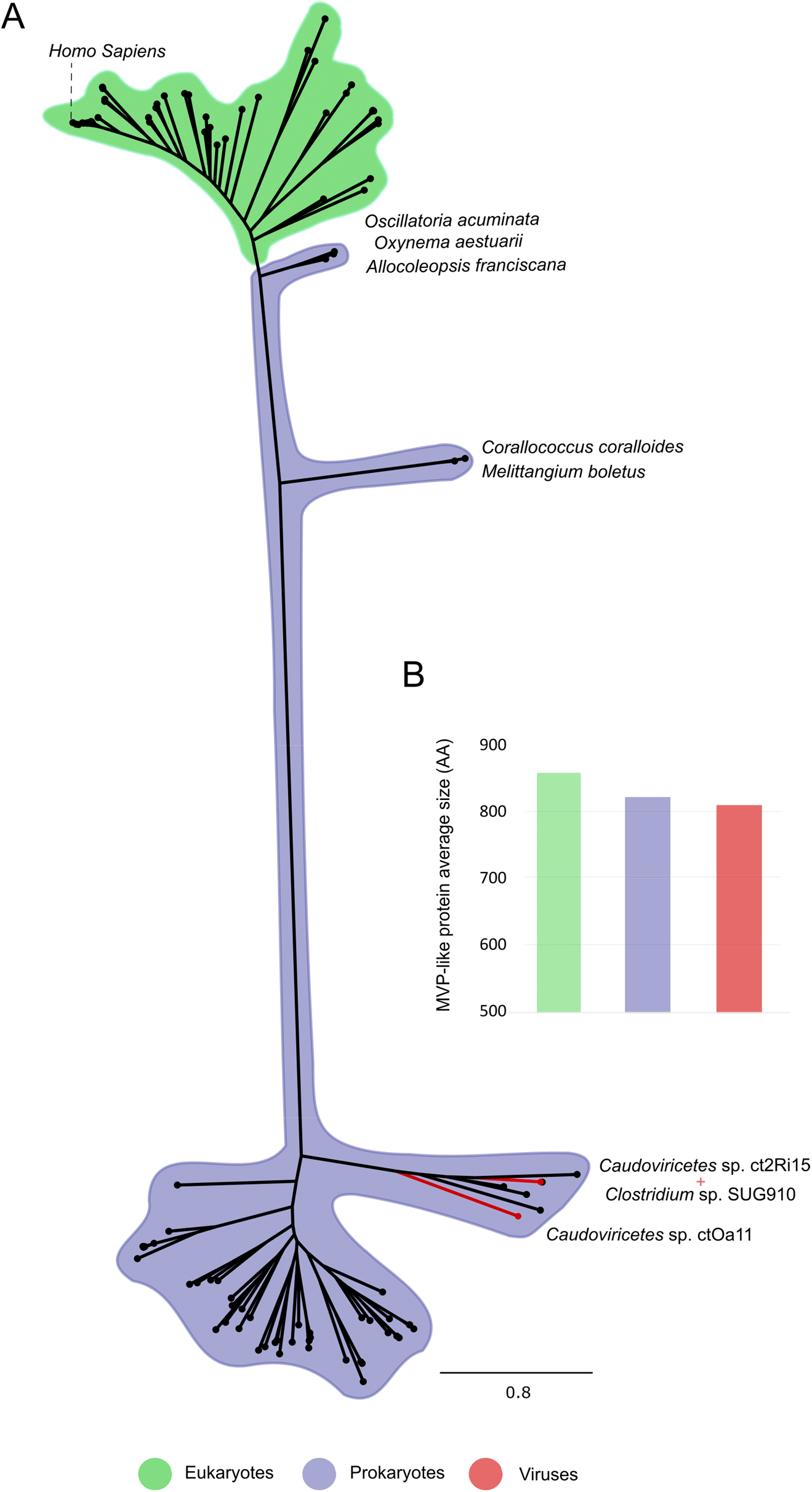
Evolutionary relationship and diversity of MVPs from selected eukaryotes, prokaryotes, and viruses. **A)** Unrooted radial consensus tree of MVPs sequences obtained by the Maximum Likelihood method. For improved visualization, species names have been omitted, but representative species names are selectively displayed. Clades are highlighted according to taxonomic group. The complete phylogenetic tree is available in **Supplemental Figure S4**. **B)** Average MVP length for each group.

### 3.4 Structural analysis

We used the AlphaFold2 system to predict the tridimensional structure of putative viral MVPs and evaluate whether the amino acid sequences would fold into protein structures comparable to *bona-fide* MVPs. Upon visual inspection, we observed an extreme resemblance to eukaryotic MVPs, including the cap-helix domain, shoulder domain, and N-terminal repeats. To verify this observation, we compared the modelled structures to the resolved structure of the human MVP using the RMSD approach. However, RMSD analysis revealed that the phage MVPs had lower values when aligned to the bacterial homologs and higher values when compared to the human protein (**Fig. 4**). For instance, the *Clostridium* sp. MVP protein had an RMSD of 0.49 when aligned to the ct2Ri15 MVP as anticipated, due the high protein sequence similarity (∼99%). However, regions of low pLDDT, represented by unstructured regions, can affect protein alignment, which can make full protein alignment challenging. Additionally, eukaryotic MVPs have an unstructured region at the C-terminal region, and the flexibility of the cap-helix domain can generate higher RMSDs.

**Fig. 4.**
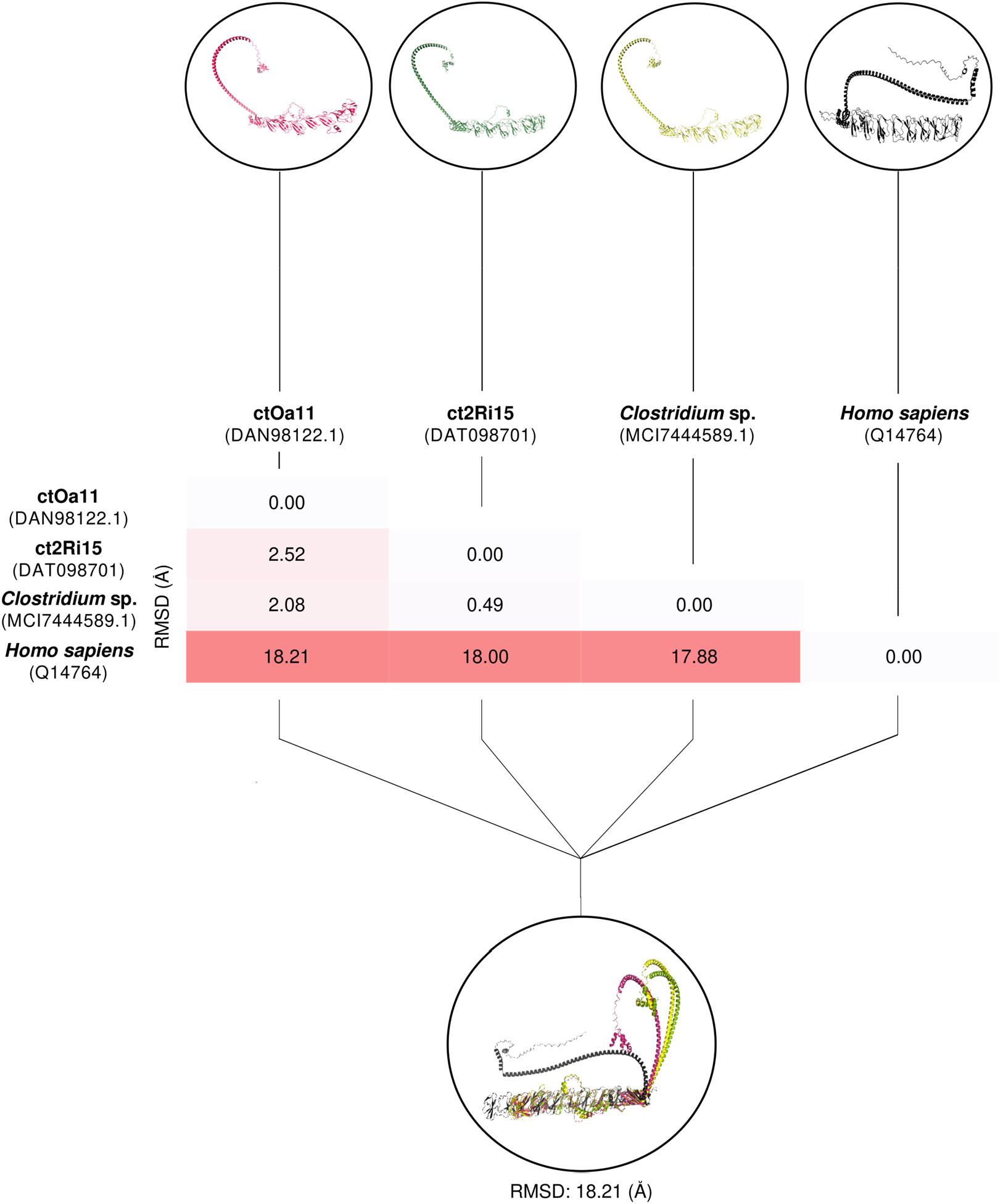
Structural comparison of predicted MVP tridimensional models. Values represent calculated RMSD from pairwise protein alignment, which indicates the level of structural difference between the predicted models. Models were generated using AlphaFold2 Colab. Protein identifiers are displayed in parentheses. RMSD, Root Mean Square Deviation.

## 4. DISCUSSION

Vaults are the largest cytoplasmic ribonucleoprotein complexes described to date and are widely distributed across diverse organisms. However, their precise function is still obscure albeit the experimental studies performed to elucidate this question. The main protein in vaults, MVP, forms the outer shell of the barrel-shaped complex structure and has not been described in viruses until now. The evolutionary dynamics of MVP are still a topic of interest for researchers. Phylogenetics reconstruction studies suggest that MVPs were likely present in the Last Eukaryotic Common Ancestor (LECA) (Daly et al., 2013b). However, the presence of MVPs homologs in species of Cyanobacteria and chemotrophic Bacteria suggests that their origin in the Last Universal Common Ancestor (LUCA) cannot be discarded (Daly et al., 2013a; Sokolskyi, 2019). Horizontal gene transfer may have been an important factor in MVPs distribution (Daly et al., 2013a, 2013b; Sokolskyi, 2019), although the direction of transfer is not yet a consensus, whether via bacteria to early eukaryotes or vice-versa.

Our study has, for the first time, revealed the existence of viral *MVP* gene homologs in two unidentified bacterial viruses from the human gut—isolates ctOa11 and ct2Ri15—from the Class *Caudoviricetes*. Current knowledge about bacteriophage taxonomy and the limited cataloged diversity, which remains significantly lower than the probable number of existing species, hindered a more precise taxonomic classification of the bacteriophages under study. Nevertheless, based on the nucleotide and protein content, we observed that ctOa11 and ct2Ri15 share a close genetic proximity.

Many *Caudoviricetes* are known to infect gut bacteria, and, as evidenced by metagenome studies, they are among the most prevalent and diverse groups of viruses in the human gut (Gulyaeva et al., 2022; Y. Zhang & Wang, 2023). The gut microbiome is a highly complex environment inhabited by a myriad of microorganisms living dynamically. Here, microbe-host and microbe-microbe interactions occur frequently (Borodovich et al., 2022; Shterzer & Mizrahi, 2015; Soucy et al., 2015). Temperate phages constantly integrate themselves into the genomes of their bacterial host as part of their life cycle. Consequently, newly sequenced bacterial species often contain prophage sequences, especially those that are not curated (Dahlman et al., 2023; Sutcliffe et al., 2023).

Based on our analyses, constrained by the current understanding of bacteriophages, the new *Caudoviricetes* phage ct2Ri15 or a closely related species may integrate itself as prophage within the genome of a *Clostridium* species. This hypothesis is based on the notable similarity between the MVPs of ct2Ri15 and *Clostridium* sp. (99.1%), the significant genetic homology between ct2Ri15 and a segment of the *Clostridium* sp. genome, the prediction of a prophage region near the *MVP* gene, and the fact that *Clostridium* is a common genus of gut bacteria. Taken together, the results of the lifestyle prediction analyses suggest that both phages are lysogenic, although one of the tools suggested that ct2Ri15 is temperate. However, given the likelihood that we are dealing with new viruses, this result must be interpreted cautiously. In the future, as more bacterial and bacteriophages genomes are being sequenced, we may discover the possibility that other MVP homologs found in some of the bacterial genomes in this work are also part of viral entities integrated within the bacterial genomes as prophages.

Until further notice, it appears that the *MVP* from ctOa11 is likely a component of that virus genome, despite sharing a significant amino acid similarity of approximately 70% with the MVP from an unidentified bacterial species in the *Lachnospiraceae* family. Both bacterial families, *Clostridiaceae* and *Lachnospiraceae*, belong to Order Eubacteriales in Phylum *Baciliota* (*Firmicutes*) and have representatives commonly found in the human gut. Previously identified bacterial MVP proteins are mostly found in genomes of heterotrophic species with gliding motility (SHAIK, 2013; Sokolskyi, 2019), which is a type of flagella-independent motility. Here, the same observation appears to remain true, where most of bacteria present in our phylogenetic reconstruction can move by this mechanism. Gliding heterotrophic bacteria need to prey on other bacteria or protists to fulfill their amino acid requirements due to incomplete biosynthetic pathways (SHAIK, 2013).

We have demonstrated the sequence similarity between viral and eukaryotic MVPs and the conservation of the classic MVP tridimensional structure. Despite the high RMSD values for viral-eukaryotic protein alignment, probably due to non-structural regions, closer visual inspection revealed a high degree of comparability between the proteins. Additionally, we noticed that, in terms of amino acid number, eukaryotic MVPs are generally longer than prokaryotic and viral counterparts, aligning with the hypothesis of an evolutionary trend toward increased protein size in eukaryotic organisms (Brocchieri & Karlin, 2005; J. Zhang, 2000). We have also validated the *MVP* genes of ctOa11 and ct2Ri15 by aligning the sequence reads to the assembled genomes. The lack of gaps and disjunctions at the edges of the genes indicates the correct assignment of the *MVP* gene in the viral genomes and rules out contamination as the origin of these genes.

The presence of *MVP* genes in phage genomes suggests an unknown role in phage infection or virion structure. One hypothesis is that the viral MVPs might function as Auxiliary Metabolic Genes (AMGs). AMGs are phage genes derived from bacteria that modulate microbial metabolism such as biosynthesis processes and cell respiration, ultimately improving viral fitness during infection (Breitbart et al., 2007). The presence of *MVP*-homologs in bacteria can be correlated with the loss of genes responsible for amino acid biosynthesis and nitrogen fixation to compensate for nitrogen deficiency, serving as an amino acid storage device (SHAIK, 2013). The bacterial species placed within the same clade as the phages in the MVP phylogenetic tree, currently the most probable host candidates, also seem to lack key nitrogen fixating genes (*nif*), as a quick BLASTp search can demonstrate. The phages could be enhancing the amino acid storage strategy by increasing MVP production. However, if the *MVP* gene were truly augmenting the fitness of the viral community, we would anticipate a rapid diffusion of this gene within the microbial community. To expand on this hypothesis to further point out the metabolic role of the viral *MVP* genes during bacterial infection, the complete function of the *MVP* gene and the vault complex must be first fully deciphered. Whether the viral MVPs play a sole metabolic role or they oligomerize to perform one of vault’s currently described functions in eukaryotes such as nutrient sequester, cargo shuttle or a yet-to-be-unveiled role, is yet to be determined.

In the study from which *Caudoviricetes* sp. ctOa11 and ct2Ri15 were sequenced (Tisza et al., 2020), non-canonical structural proteins, i.e., not the usual capsid proteins, were computationally detected in these viral genomes. They were further confirmed through lab culture experiments to form virus-like shells, thus raising the possibility of an unconventional scenario where the phage MVPs can oligomerize to form viral shells in the form of vaults. This hypothesis is considered less plausible due to the presence of phage structural proteins, such as the major capsid protein, in the genomes of both viruses. Additionally, the localization of phage *MVPs* outside the structural genomic clusters, a common characteristic of bacteriophage genomes, further reduces the likelihood of this scenario. Despite the absence of viruses with radially symmetrical halves joined together like the vault particle, the structural and biochemical characteristics of such structures, such as the presence of numerous copies of a single polypeptide chain, self-assembly, metastability, hollow interior and affinity for nucleic acids, support their potential use as shells by viral replicators. These observations also lend support to the fascinating hypothesis that vaults could be remnants of ancient viral symbionts of eukaryotic cells (Nandy et al., 2009). If validated, this theory would have significant implications for current models of the evolution of viruses and their host organisms.

Recent developments in genome sequencing and the growing number of newly sequenced microorganisms are shedding light on the genome diversity of viruses and bacteria. Described here for the first time, the identification of two *MVP* homologs in the genome of new viral entities from the “viral dark matter” offers new insights into the phylogenetic distribution of this protein. In this research, we computationally characterized two atypical viruses, which are likely to be taxonomically related. While only two MVP-bearing viruses have been identified so far, it is possible that more exist and additional metagenomic research should be conducted to uncover them. Finally, any conclusion to be drawn concerning the role of MVP in phages would require first the elucidation of MVPs function in bacteria. More insights would be gained through additional experimental studies on the neglected vault complex and MVPs, particularly the bacterial homologs, to decode their function outside Eukarya. Likewise, *in vivo* and *in vitro* studies on these unique viruses would be beneficial in unraveling their structure and pathogenesis.

## Author Contributions

Thiago Mendonça dos Santos and Luiz Eduardo Vieira Del Bem contributed to the study conception and design. Material preparation and data collection were performed by Thiago Mendonça dos Santos. Analyses were performed by Thiago Mendonça dos Santos, Emerson Galvão Colaço and Helena Ferreira Gomes. The first draft of the manuscript was written by Thiago Mendonça dos Santos. Luiz Eduardo Vieira Del Bem supervised the study and revised the manuscript. All authors read and approved the final manuscript.

