## Supplemental Figure S1 for "Characterization and Phylogenetic Insights into the First Viral Major Vault Proteins (MVPs), Identified in Bacteriophages from the Human Gut"

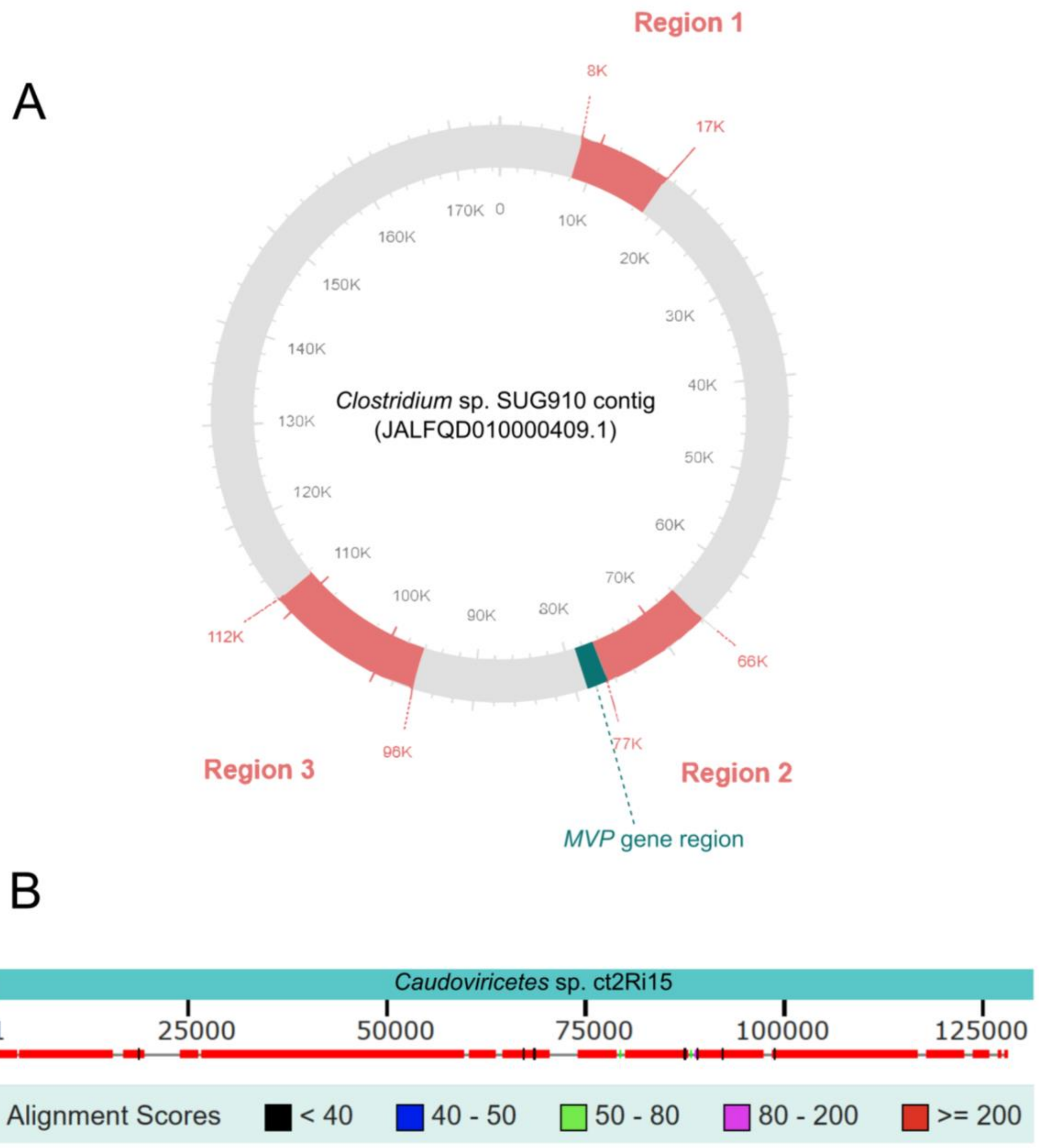


**Supplemental Figure S1.** Prophage search results. **A)** Genomic map of *Clostridium* sp. SUG910 contig highlighting predicted regions of prophages integrated to the bacterial genome. Regions are predicted by PHASTER web-server. Red color denotes incomplete prophage regions. **B)** Graphical representation of BLASTn results between *Caudoviricetes* sp. ct2Ri15 genome and *Clostridium* sp. SUG910 contig. Red boxes indicate high-score alignments of phage genome to bacterial contig.
