## Supplemental Figure S2 for "Characterization and Phylogenetic Insights into the First Viral Major Vault Proteins (MVPs), Identified in Bacteriophages from the Human Gut"

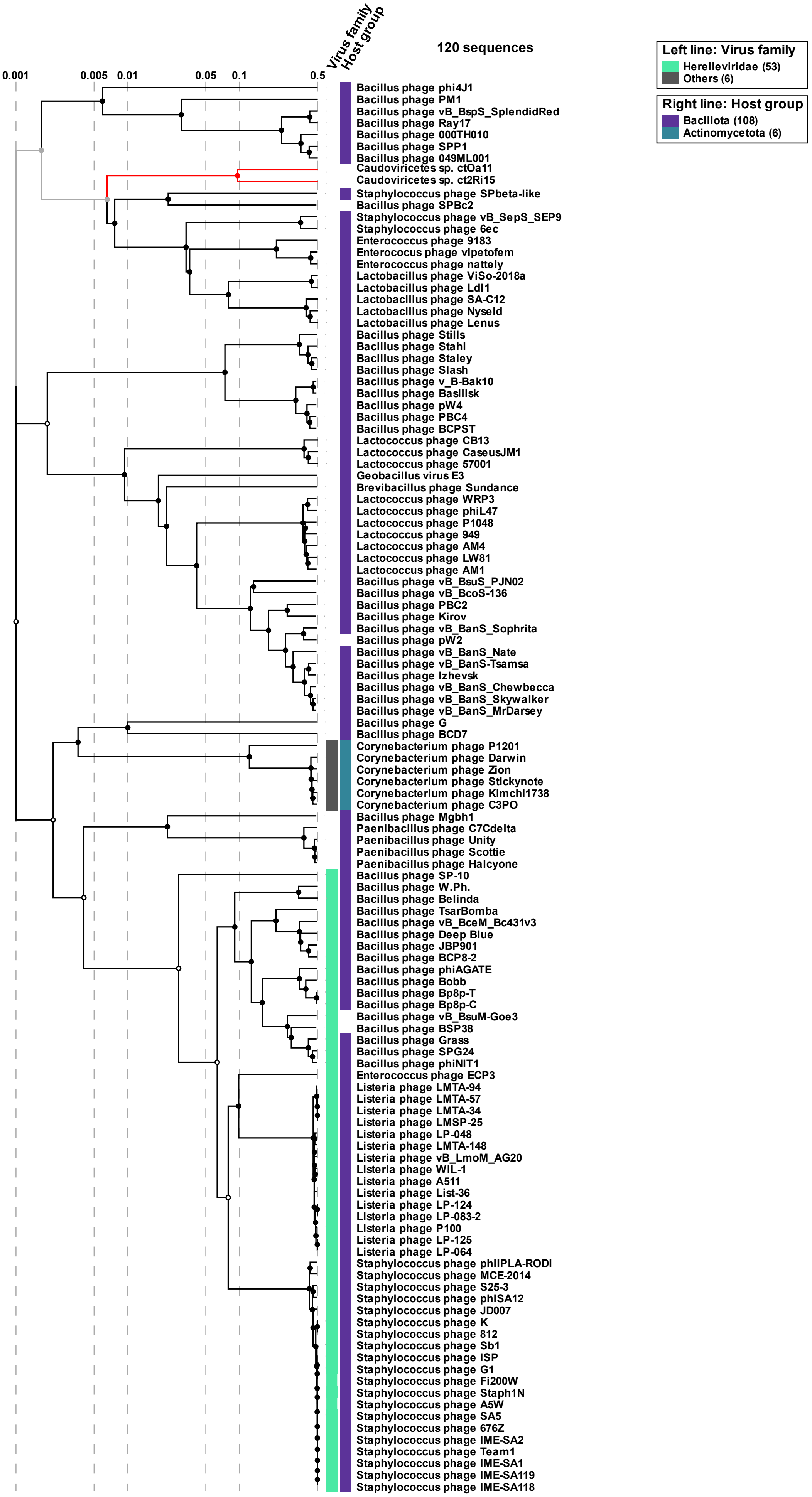

120 sequences
Virus family
Host group
0.001
0.005
0.01
0.05
0.1
0.5
Left line: Virus family
Herelleviridae (53)
Others (6)
Right line: Host group
Bacillota (108)
Actinomycetota (6)
Bacillus phage phi4J1
Bacillus phage PM1
Bacillus phage vB_BspS_SplendidRed
Bacillus phage Ray17
Bacillus phage 000TH010
Bacillus phage SPP1
Bacillus phage 049ML001
Caudoviricetes sp. ctOa11
Caudoviricetes sp. ct2Ri15
Staphylococcus phage SPbeta-like
Bacillus phage SPBc2
Staphylococcus phage vB_SepS_SEP9
Staphylococcus phage 6ec
Enterococcus phage 9183
Enterococcus phage vipetofem
Enterococcus phage nattely
Lactobacillus phage ViSo-2018a
Lactobacillus phage Ldl1
Lactobacillus phage SA-C12
Lactobacillus phage Nyseid
Lactobacillus phage Lenus
Bacillus phage Stills
Bacillus phage Stahl
Bacillus phage Staley
Bacillus phage Slash
Bacillus phage v_B-Bak10
Bacillus phage Basilisk
Bacillus phage pW4
Bacillus phage PBC4
Bacillus phage BCPST
Lactococcus phage CB13
Lactococcus phage CaseusJM1
Lactococcus phage 57001
Geobacillus virus E3
Brevibacillus phage Sundance
Lactococcus phage WRP3
Lactococcus phage phiL47
Lactococcus phage P1048
Lactococcus phage 949
Lactococcus phage AM4
Lactococcus phage LW81
Lactococcus phage AM1
Bacillus phage vB_BsuS_PJN02
Bacillus phage vB_BcoS-136
Bacillus phage PBC2
Bacillus phage Kirov
Bacillus phage vB_BanS_Sophrita
Bacillus phage pW2
Bacillus phage vB_BanS_Nate
Bacillus phage vB_BanS-Tsamsa
Bacillus phage Izhevsk
Bacillus phage vB_BanS_Chewbecca
Bacillus phage vB_BanS_Skywalker
Bacillus phage vB_BanS_MrDarsey
Bacillus phage G
Bacillus phage BCD7
Corynebacterium phage P1201
Corynebacterium phage Darwin
Corynebacterium phage Zion
Corynebacterium phage Stickynote
Corynebacterium phage Kimchi1738
Corynebacterium phage C3PO
Bacillus phage Mgbh1
Paenibacillus phage C7Cdelta
Paenibacillus phage Unity
Paenibacillus phage Scottie
Paenibacillus phage Halcyone
Bacillus phage SP-10
Bacillus phage W.Ph.
Bacillus phage Belinda
Bacillus phage TsarBomba
Bacillus phage vB_BceM_Bc431v3
Bacillus phage Deep Blue
Bacillus phage JBP901
Bacillus phage BCP8-2
Bacillus phage phiAGATE
Bacillus phage Bobb
Bacillus phage Bp8p-T
Bacillus phage Bp8p-C
Bacillus phage vB_BsuM-Goe3
Bacillus phage BSP38
Bacillus phage Grass
Bacillus phage SPG24
Bacillus phage phiNIT1
Enterococcus phage ECP3
Listeria phage LMTA-94
Listeria phage LMTA-57
Listeria phage LMTA-34
Listeria phage LMSP-25
Listeria phage LP-048
Listeria phage LMTA-148
Listeria phage vB_LmoM_AG20
Listeria phage WIL-1
Listeria phage A511
Listeria phage List-36
Listeria phage LP-124
Listeria phage LP-083-2
Listeria phage P100
Listeria phage LP-125
Listeria phage LP-064
Staphylococcus phage philPLA-RODI
Staphylococcus phage MCE-2014
Staphylococcus phage S25-3
Staphylococcus phage phiSA12
Staphylococcus phage JD007
Staphylococcus phage K
Staphylococcus phage 812
Staphylococcus phage Sb1
Staphylococcus phage ISP
Staphylococcus phage G1
Staphylococcus phage Fi200W
Staphylococcus phage Staph1N
Staphylococcus phage A5W
Staphylococcus phage SA5
Staphylococcus phage 676Z
Staphylococcus phage IME-SA2
Staphylococcus phage Team1
Staphylococcus phage IME-SA1
Staphylococcus phage IME-SA119
Staphylococcus phage IME-SA118

**Supplemental Figure S2.** Complete proteomic tree generated by VipTree based on genome-wide similarities as determined by tBLASTx plotted on a log scale. The analysis includes genome sequences of *Caudoviricetes* sp. ct2Ri15, ctOa11, along with genome sequences of phages from Virus-Host DB (RefSeq release 219). Right and left lines are color-coded according to host group and virus family, respectively. For reference, ct2Ri15 and ctOa11 phages are highlighted in red.
