## Supplemental Figure S3 for "Characterization and Phylogenetic Insights into the First Viral Major Vault Proteins (MVPs), Identified in Bacteriophages from the Human Gut"

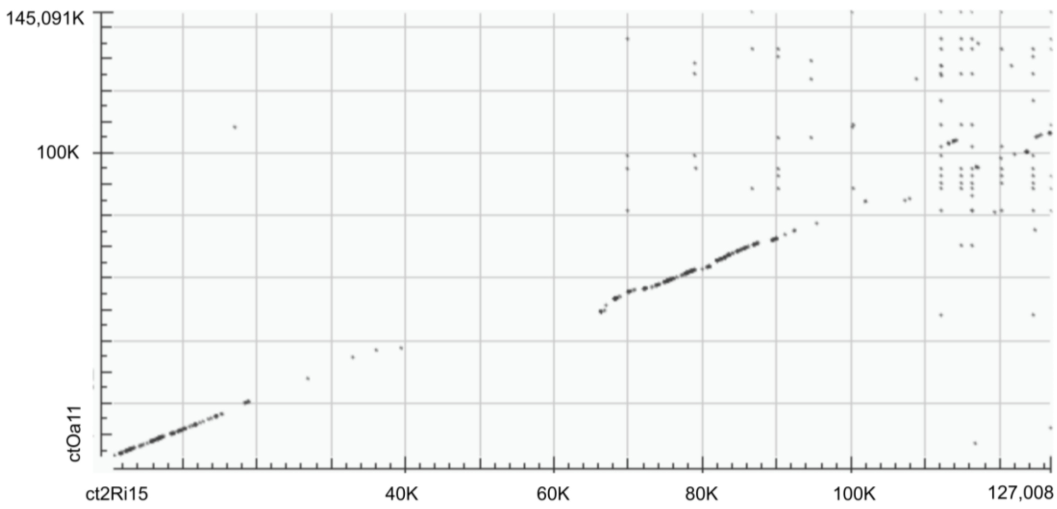


**Supplemental Figure S3.** BLASTn dotplot output displaying the alignment of *Caudoviricetes* sp. ct2Ri15 and ctOa11 genome sequences. Dots indicates the location of BLASTn hits (e-value < $1.10^{-05}$).
