## Supplemental Figure S4 for "Characterization and Phylogenetic Insights into the First Viral Major Vault Proteins (MVPs), Identified in Bacteriophages from the Human Gut"

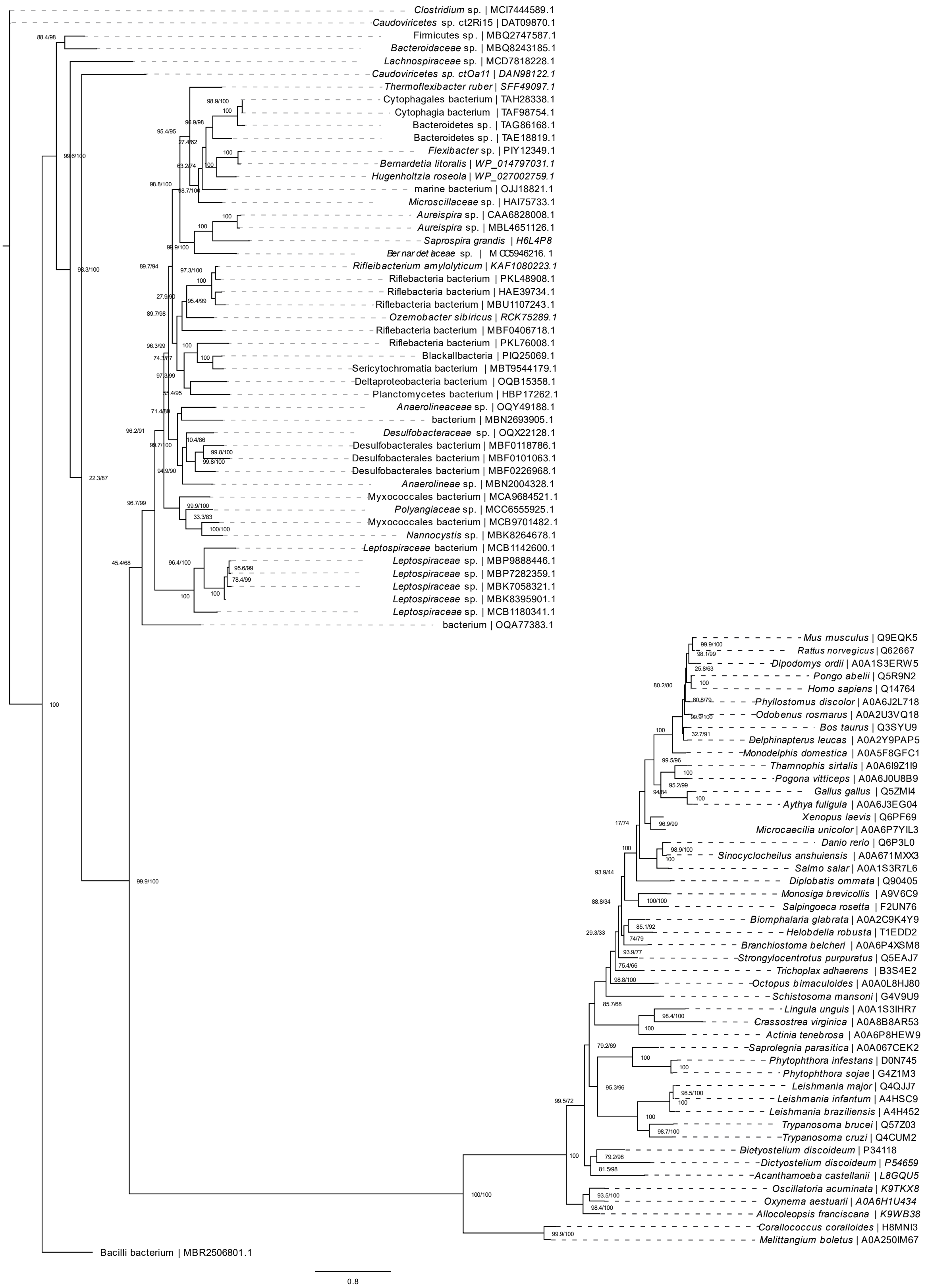
Clostridium sp. | MCI7444589.1
Caudoviricetes sp. ct2Ri15 | DAT09870.1
Firmicutes sp. | MBQ2747587.1
Bacteroidaceae sp. | MBQ8243185.1
Lachnospiraceae sp. | MCD7818228.1
Caudoviricetes sp. ctOa11 | DAN98122.1
Thermoflexibacter ruber | SFF49097.1
Cytophagales bacterium | TAH28338.1
Cytophagia bacterium | TAF98754.1
Bacteroidetes sp. | TAG86168.1
Bacteroidetes sp. | TAE18819.1
Flexibacter sp. | PIY12349.1
Bernardetia litoralis | WP_014797031.1
Hugenholtzia roseola | WP_027002759.1
marine bacterium | OJJ18821.1
Microscillaceae sp. | HAI75733.1
Aureispira sp. | CAA6828008.1
Aureispira sp. | MBL4651126.1
Saprospira grandis | H6L4P8
Ber nar det iaceae sp. | MCC5946216.1
Rifleibacterium amylolyticum | KAF1080223.1
Riflebacteria bacterium | PKL48908.1
Riflebacteria bacterium | HAE39734.1
Riflebacteria bacterium | MBU1107243.1
Ozemobacter sibiricus | RCK75289.1
Riflebacteria bacterium | MBF0406718.1
Riflebacteria bacterium | PKL76008.1
Blackallbacteria | PIQ25069.1
Sericytochromatia bacterium | MBT9544179.1
Deltaproteobacteria bacterium | OQB15358.1
Planctomycetes bacterium | HBP17262.1
Anaerolineaceae sp. | OQY49188.1
bacterium | MBN2693905.1
Desulfobacteraceae sp. | OQX22128.1
Desulfobacterales bacterium | MBF0118786.1
Desulfobacterales bacterium | MBF0101063.1
Desulfobacterales bacterium | MBF0226968.1
Anaerolineae sp. | MBN2004328.1
Myxococcales bacterium | MCA9684521.1
Polyangiaceae sp. | MCC6555925.1
Myxococcales bacterium | MCB9701482.1
Nannocystis sp. | MBK8264678.1
Leptospiraceae bacterium | MCB1142600.1
Leptospiraceae sp. | MBP9888446.1
Leptospiraceae sp. | MBP7282359.1
Leptospiraceae sp. | MBK7058321.1
Leptospiraceae sp. | MBK8395901.1
Leptospiraceae sp. | MCB1180341.1
bacterium | OQA77383.1
Mus musculus | Q9EQK5
Rattus norvegicus | Q62667
Dipodomys ordii | A0A1S3ERW5
Pongo abelii | Q5R9N2
Homo sapiens | Q14764
Phyllostomus discolor | A0A6J2L718
Odobenus rosmarus | A0A2U3VQ18
Bos taurus | Q3SYU9
Delphinapterus leucas | A0A2Y9PAP5
Monodelphis domestica | A0A5F8GFC1
Thamnophis sirtalis | A0A6I9Z1I9
Pogona vitticeps | A0A6J0U8B9
Gallus gallus | Q5ZMI4
Aythya fuligula | A0A6J3EG04
Xenopus laevis | Q6PF69
Microcaecilia unicolor | A0A6P7YIL3
Danio rerio | Q6P3L0
Sinocyclocheilus anshuiensis | A0A671MXX3
Salmo salar | A0A1S3R7L6
Diplobatis ommata | Q90405
Monosiga brevicollis | A9V6C9
Salpingoeca rosetta | F2UN76
Biomphalaria glabrata | A0A2C9K4Y9
Helobdella robusta | T1EDD2
Branchiostoma belcheri | A0A6P4XSM8
Strongylocentrotus purpuratus | Q5EAJ7
Trichoplax adhaerens | B3S4E2
Octopus bimaculoides | A0A0L8HJ80
Schistosoma mansoni | G4V9U9
Lingula unguis | A0A1S3IHR7
Crassostrea virginica | A0A8B8AR53
Actinia tenebrosa | A0A6P8HEW9
Saprolegnia parasitica | A0A067CEK2
Phytophthora infestans | D0N745
Phytophthora sojae | G4Z1M3
Leishmania major | Q4QJJ7
Leishmania infantum | A4HSC9
Leishmania braziliensis | A4H452
Trypanosoma brucei | Q57Z03
Trypanosoma cruzi | Q4CUM2
Dictyostelium discoideum | P34118
Dictyostelium discoideum | P54659
Acanthamoeba castellanii | L8GQU5
Oscillatoria acuminata | K9TKX8
Oxynema aestuarii | A0A6H1U434
Allocoleopsis franciscana | K9WB38
Corallococcus coralloides | H8MNI3
Melittangium boletus | A0A250IM67
Bacilli bacterium | MBR2506801.1
0.8

**Supplemental Figure S4.** Unrooted radial consensus tree of 98 MVPs sequences. The tree was generated by IQ-TREE through the Maximum Likelihood method and branch support values are displayed on each node. Numbers in parentheses are SH-aLRT support (%) / ultrafast bootstrap support (%). If value is the same for both, only one number is displayed. Protein identifiers are displayed after the vertical bars. Details on the parameters used in generating this phylogenetic tree are outlined in the Methods section.
